# Human activities drive novel behaviours and transitions in dolphins

**DOI:** 10.1101/2025.07.10.663819

**Authors:** Imran Samad, Harshal Patil, Mauricio Cantor, Damien Farine, Dipani Sutaria, Kartik Shanker

## Abstract

Intensifying human activities are reshaping coastal ecosystems, yet their impact on wildlife behaviour and survival remain poorly understood. We conducted drone-based focal-group follows of endangered Indian Ocean humpback dolphin groups in Goa, India, to quantify how tourism and fisheries jointly impact behavioural states and transitions. By integrating machine learning and georeferencing techniques, we found that both human activities triggered distinct behavioural transitions absent in undisturbed groups. Dolphins expressed ‘escape’ behaviours exclusively near tourist boats and were more likely to transition into foraging near fishing nets (mostly purse-seines). Smaller groups reacted more strongly to tourist boats and interacted more frequently with fishing nets, especially in seasons when fish landing data showed a decline in their prey. These findings offer new insights into the behavioural mechanisms underlying the co-occurrence of coastal dolphins with humans and provide broader guidelines for managing tourism and fisheries to reduce anthropogenic pressure on threatened delphinids.

## Introduction

Human activities are increasingly forcing wildlife to adapt to landscapes altered by unnatural resource extraction and habitat modification (Allan et al., 2019). Anthropogenic pressures often drive direct wildlife mortality and demographic shifts (Tuomainen & Candolin, 2011). Even non-lethal human activities can translate to changes in animal behaviour and population dynamics (Smith et al., 2024), which is especially detrimental for range-restricted species (Chen et al., 2023). However, our understanding of how multiple, co-occurring human activities shape the behavioural dynamics of animal groups remains limited.

Coastal cetaceans are widely threatened by high overlap and recurrent interactions with human activities, including fisheries and tourism (Braulik et al., 2023). Coastal fisheries deploy a range of gear (e.g., gillnets, trawl nets, purse seines) that ‘attract’ cetaceans to prey concentrations but can result in mutual harm - entanglement, gear damage and catch loss (Jog et al., 2022). Tourist boats are an additional increasing concern (e.g., Constantine, Brunton & Denis, 2004) as these can chase groups over long distances leading to stress and injury. Human activities like these can impact dolphin populations directly through bycatch and indirectly through alteration of their natural behaviour and habitat-use (Piwetz, Lundquist & Würsig, 2015; Sumanapala & Wolf 2024).

Cetacean behaviour may be particularly impacted when these disturbances are active simultaneously, even though fisheries and tourism are important sources of livelihoods in many regions (FAO, 2022; Sumanapala & Wolf 2024). Thus, managing coastal dolphin populations requires detailed information on how and when dolphin groups react to these activities – both independently and jointly – especially as the effects may not simply be additive. For instance, dolphins can often be attracted to fishing vessels due to foraging opportunities, but how do they engage with fisheries in the presence of tourist boats?

The effects of disturbances on dolphins are typically studied using a control-impact design, where changes in the distribution of behavioural states are attributed to the presence of the impact (Constantine, Brunton & Denis, 2004; Jog et al., 2024). However, behavioural states are only part of the equation: to be in a behavioural state, an individual must transition from another. In nature, many behavioural transitions are relatively conserved (Minasandra et al., 2025). For example, an individual needs to repeatedly transition between searching and foraging states to acquire the energy to survive (Charnov 1976). While many studies address the effect of various external factors on behavioural states, few address the factors governing behavioural transitions (Gundermann et al., 2023). This is important because while individuals may express the same overall distribution of behaviours across disturbances, there may be novel behavioural state transitions that may not otherwise exist under natural conditions – a cryptic form of behavioural change.

One challenge with capturing the distinct effects of human activities on dolphin behaviour is the need to track groups at high resolutions and record both behavioural states and their transitions. These data then need to be replicated under different contexts in natural settings. Drones have emerged as an accessible solution for capturing replicated events of cetacean behaviour (Hartman, van der Harst & Vilela, 2020). Combined with novel analytical tools for automatically detecting and tracking individuals (e.g., Koger et al., 2022; Samad et al., 2025), drone-based research is revolutionizing the study of dolphin behaviour, offering unprecedented precision in detecting behavioural transitions in responses to human activities.

Here, we conducted drone-based focal group follows of Indian Ocean humpback dolphins (IOHD; *Sousa plumbea*) in Goa on the west coast of India, to quantify their fine-scale responses to co-occurring fishing nets and tourist vessels. This endangered coastal cetacean inhabits a narrow band (<5 km from the shore) along India’s west coast where the central-west region is particularly important (Samad, Sutaria & Shanker, 2024). In Goa, IOHD are frequently observed interacting with fisheries and tourism (Sanjeev et al., 2016) but we lack empirical information to develop guidelines for managing their interactions with dolphins. Our objective was to quantify how and when dolphin groups react to fishing and tourism activities, both independently and when they co-occur. We first classify behavioural states from drone footage to estimate behavioural transition probabilities across contexts – none, fishing, tourism, or both combined – and then quantify how behavioural impacts vary with dolphin group size and environmental factors. We conclude by discussing how fisheries and tourism management can contribute to conserving this endangered dolphin species.

## 2. Methods

### 2.1. Study site

The central west coast of India is an important marine mammal area (IUCN, 2025) that supports high densities of Indian Ocean humpback dolphins (Sutaria & Jefferson, 2005). In Goa, several rivers flow into the sea forming productive estuarine habitats that attract both dolphins and fisheries. In north Goa, such habitats overlap with areas of high dolphin-watching tourism, and humpback dolphins regularly interact with both fisheries and tourism (Sanjeev et al., 2016). We sampled areas from vantage points more than 50 meters in height in north Goa (Figure 1) from where dolphins, fishing boats and tourist boats could be observed.

**Figure 1:.**
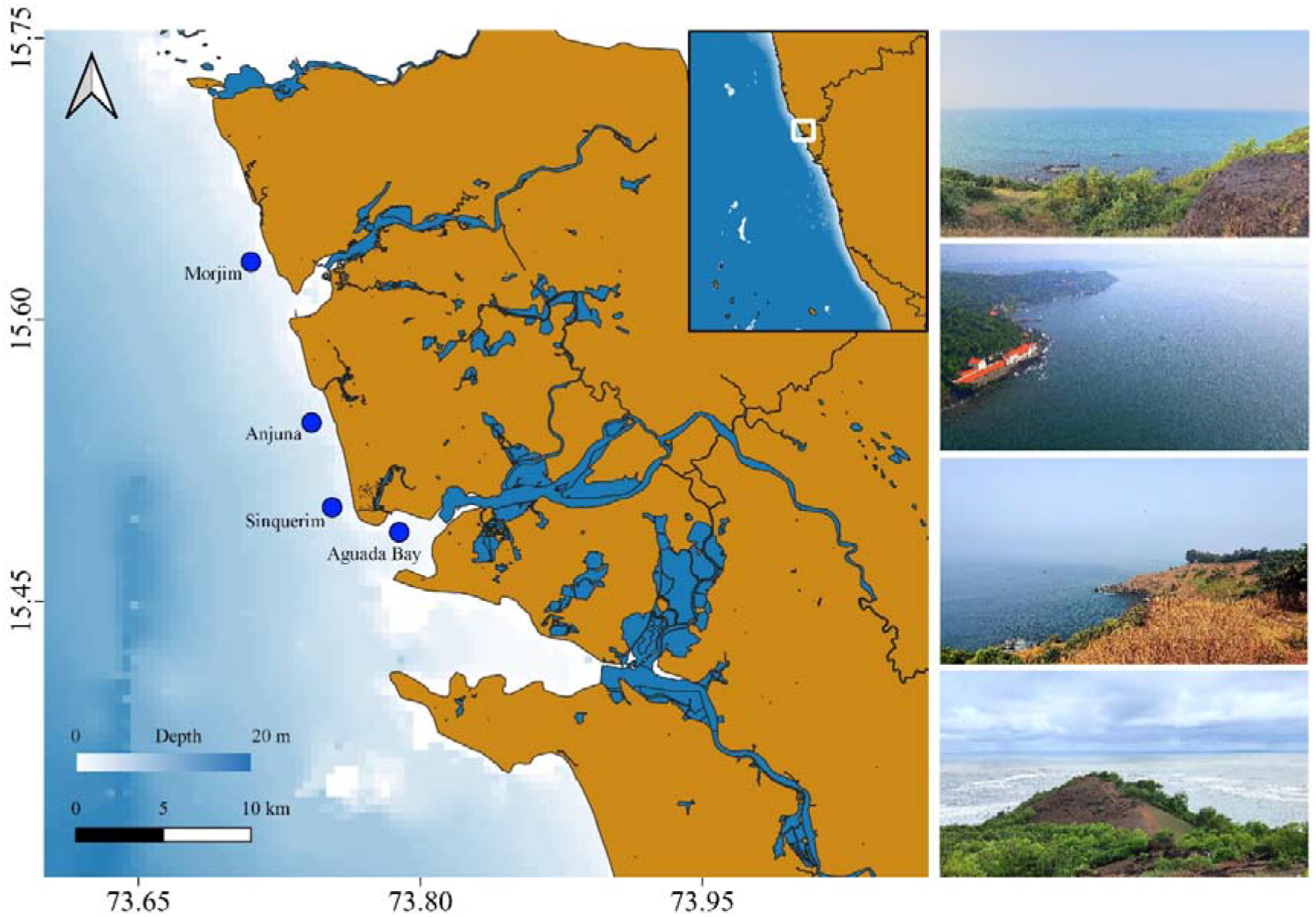
Map of the study area and sampling locations, along the central western coast of India.

### 2.2. Data collection

Between October 2023 and April 2024, 2-3 field researchers visited sampling sites 4-5 days a week for 1-3 hours in the morning and scanned the sea at regular intervals to detect dolphins (Cecchetti et al., 2018). Once dolphins were sighted, we followed them by flying a DJI Mini 3 Pro drone at a height of 60-100 meters above sea level to record their movement, group dynamics, and behaviours in the absence of, and in response to, actively fishing and tourist boats. We arbitrarily selected a dolphin group (i.e., one or more individuals within 10 meters of each other; Syme, Kiszka & Parra, 2022) and followed it until lost or drone battery dropped below 25%, then repeated the process. We continued to follow the focal group if it merged with another; if it split, we arbitrarily selected one sub-group to continue following. Occasionally the drone either recorded several groups moving close together or did not capture all group members.

Simultaneously, to understand how fish-prey availability may impact human-dolphin interactions seasonally, we visited fish landing areas close to our sampling sites to have a local estimate of prey abundance. We recorded the total weight and approximate length of fish caught by purse seines and gillnets - the most active gear in the area. Using data on the size of common fish species that dolphins prey upon (Lin et al., 2023), we estimated catch per unit effort (CPUE) as the total catch weight/fishing effort (in hours) to understand trends in fish-catch availability across months. By visually identifying sharp changes in CPUE, we broadly defined two major seasons (‘high’ and ‘low’ fish-catch) to test the extent of its impact on dolphin-fisheries and -tourism interactions.

### 2.3. Data analysis

We identified and tracked dolphin groups in drone videos using a convolutional neural network and translated their pixel-level trajectories into geo-referenced trajectories on the water surface (Samad et al., 2025). From these trajectories, we estimated group size, speed and movement direction every second (Figure 2). Instances of groups diving, merging or splitting were manually verified in the videos. Behavioural states were categorised as *travelling, socialising, foraging*, and *escape*; only dive events were categorised as *foraging* or *escape* (Table 1). Since we were interested in broad level changes in behaviour, we did not separate behavioural states further (e.g., slow vs. fast travel, milling) or consider less frequent behavioural events (e.g., breaching).

**Table 1:.** Description of the four behavioural states used in our analysis, adapted from Jog et al. (2024). Travel and socialise were estimated based on changes in the movement direction of the group, while only dives were classified as forage or escape.

| Behaviour | Description |  |
| --- | --- | --- |
|  | Jog et al. (2024) | Modified for overhead observation (this study) |
| Travel | Uniform directional movement with rhythmic resurfacing bouts | Constant movement in a fixed direction with occasional shallow dives. Change in the movement direction of the group was less than 30 degrees. |
| Socialise | High surface activity like leaping, chasing, etc. including bodily contact | Erratic movement with abrupt changes in movement direction and speed often accompanied by physical touches. Change in the movement direction of the group was more than 30 degrees. |
| Forage | Variable movement with deep dives and fish chases | Deep dives with unpredictable surfacing patterns followed by slow movement upon resurfacing. |
| Escape | NA | Sudden, unpredictable dives often distinct from dives related to travel, socialising, milling, or foraging. |

**Figure 2:.**
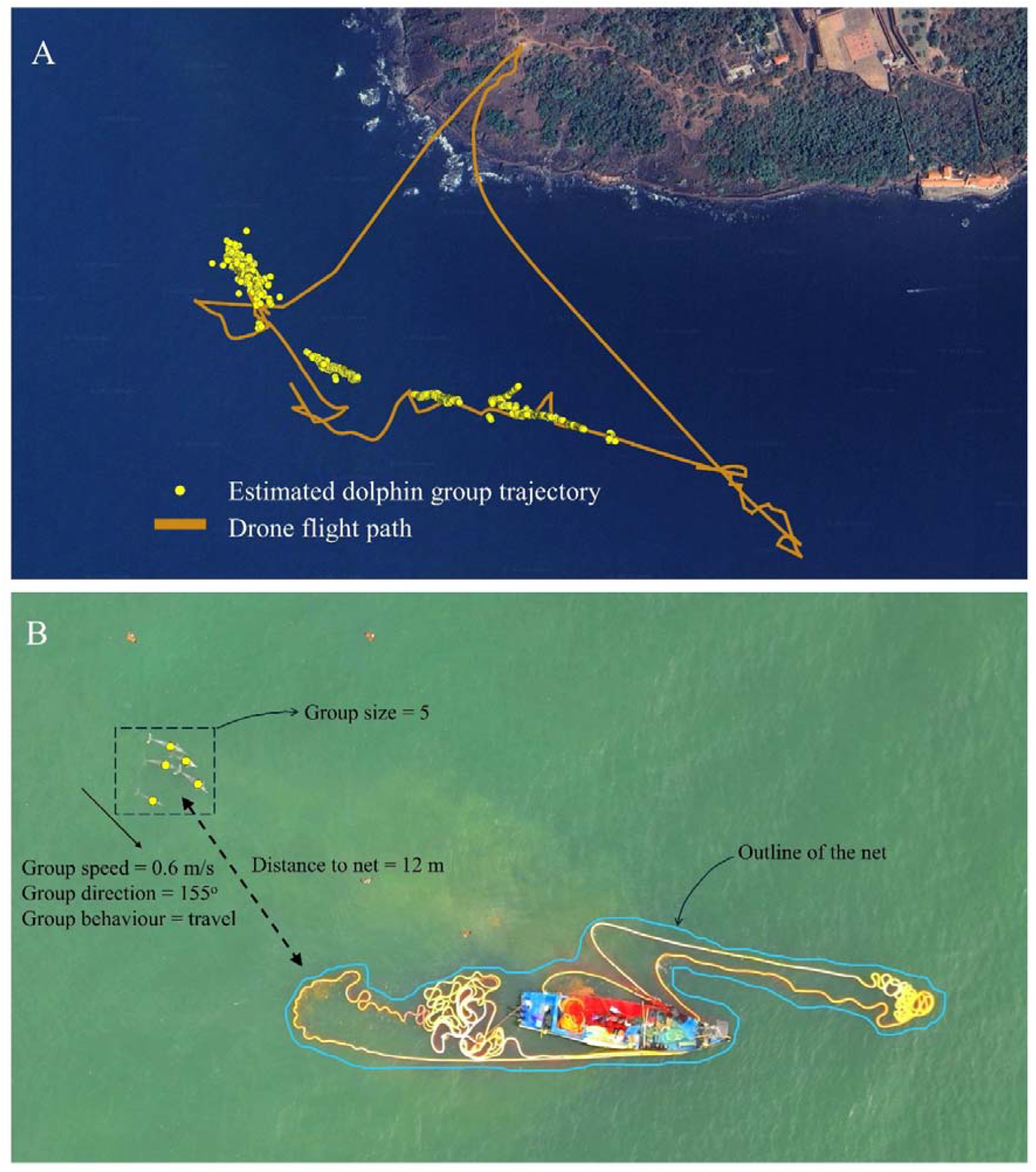
Representation of the drone flight and predicted dolphin trajectories (A) and schematic of the process of extracting data from videos (B).

We extracted data on tourist boats with a pre-trained YOLO v8 model (Varghese & Sambath, 2024). We traced out fishing nets manually in videos and extracted pixel coordinates describing their shapes (Samad et al., 2025). We geo-referenced, converted to on-ground locations, and synchronised the pixels of all human activity objects with dolphin group trajectories using video timestamps. We also extracted group-follow level data on the distance to the closest fishing net, the distance to the closest tourist boat, and the number of surrounding tourist boats (Figure 2; Supplementary Material S1).

To understand how fisheries and tourism impacted dolphin behaviour, and the factors underlying behavioural state transitions, we split our dolphin group-follow data into four pseudo-treatments: “no boats” (no tourism or fishery activities), “tourist boats” (active tourist boats nearby), “fishing nets” (fishing boats and deployed nets), and “both” (both tourist vessels and fishing boats with nets). Behavioural data for each pseudo-treatment came from an independent drone flight. For each pseudo-treatment, we estimated the probability of a dolphin group being in one of the four behavioural states (Table 1) as the mean proportion of that behavioural state across group-follows, and the probability of transitioning from one state to another during group-follows. We used multinomial regression models (Elff, 2024) in R (R Core team, 2025) to model both behavioural states and their transitions as: *Behavioural state or transition ∼ group size + neighbouring groups (present/absent) + distance to closest tourist vessel + number of surrounding tourist vessels + distance to closest fishing net + seasonal fish-catch availability (high/low)*.

## 3. Results

Over six months, we spent 104 days sampling for dolphins and recorded their presence on 77 days (Table S1, Supplementary Material S1). Our total drone flight duration was 40.4 hours in which we recorded dolphins for 22.1 hours (Table 2), all within 2 km of the coastline. Based on these observations, dolphin activity was highest in October/November (mean = 31.4 ± 10.5 SE encounters/hr), tourism activity peaked in December and dropped in March-April, and fisheries peaked in February (Figure S1, Supplementary Material S1).

**Table 2:.** A summary of sampling effort over the 22.1 hours of dolphin footage recorded. Focal group follows refer to the group that was formally being tracked on the drone while total number of group-follows include other groups that were recorded in the process.

| Pseudo-treatment | Number of focal group follows | Mean follow duration | Mean (SD) number of dolphins per group | Total number of group-follows |
| --- | --- | --- | --- | --- |
| No boats | 9 | 5m 14s | 3.4 (3.5) | 55 |
| Tourist boats | 57 | 11m 33s | 2.4 (2.7) | 477 |
| Fishing net | 13 | 13m 49s | 2.0 (1.7) | 133 |
| Both | 11 | 16m 49s | 1.8 (1.6) | 328 |

Most tourist vessels were more than 12 meters in length, used an outboard engine and carried 10-20 passengers at a time. Speedboats and large catamarans were comparatively rare but also chased dolphins. Most actively fishing vessels used purse seines to often target shrimps. Fewer vessels (<20%) used monofilament gillnets of <60 mm mesh size, and primarily target Indian mackerel (*Rastrelliger kanagurta*) and Indian oil sardine (*Sardinella longiceps*) of <30 cm in total length. CPUE was high in October/November (251 ± 48.7 kg/hr; hereby defined as high-catch season) and remained low in the remaining months (117 ± 16.8 kg/hr; low-catch season) (Figure S2, Supplementary Material S1).

### 3.1. Dolphin behavioural states and transitions

When undisturbed (pseudo-treatment “no boats”), dolphin groups predominantly travelled (probability = 0.9) followed by socialising (0.07) and foraging (0.03). When tourist boats approached (“tourist boats”), the probability of socialising or foraging decreased by 36% and 50%, respectively. In the presence of fishing nets (“fishing nets”), foraging probability increased by 330% and was highest in the low-catch season (by 37%). When tourist boats approached dolphins, their groups were 85% less likely to forage in the presence of fishing nets but they also showed lower levels of escape behaviour (“both”; Figure 3).

**Figure 3:.**
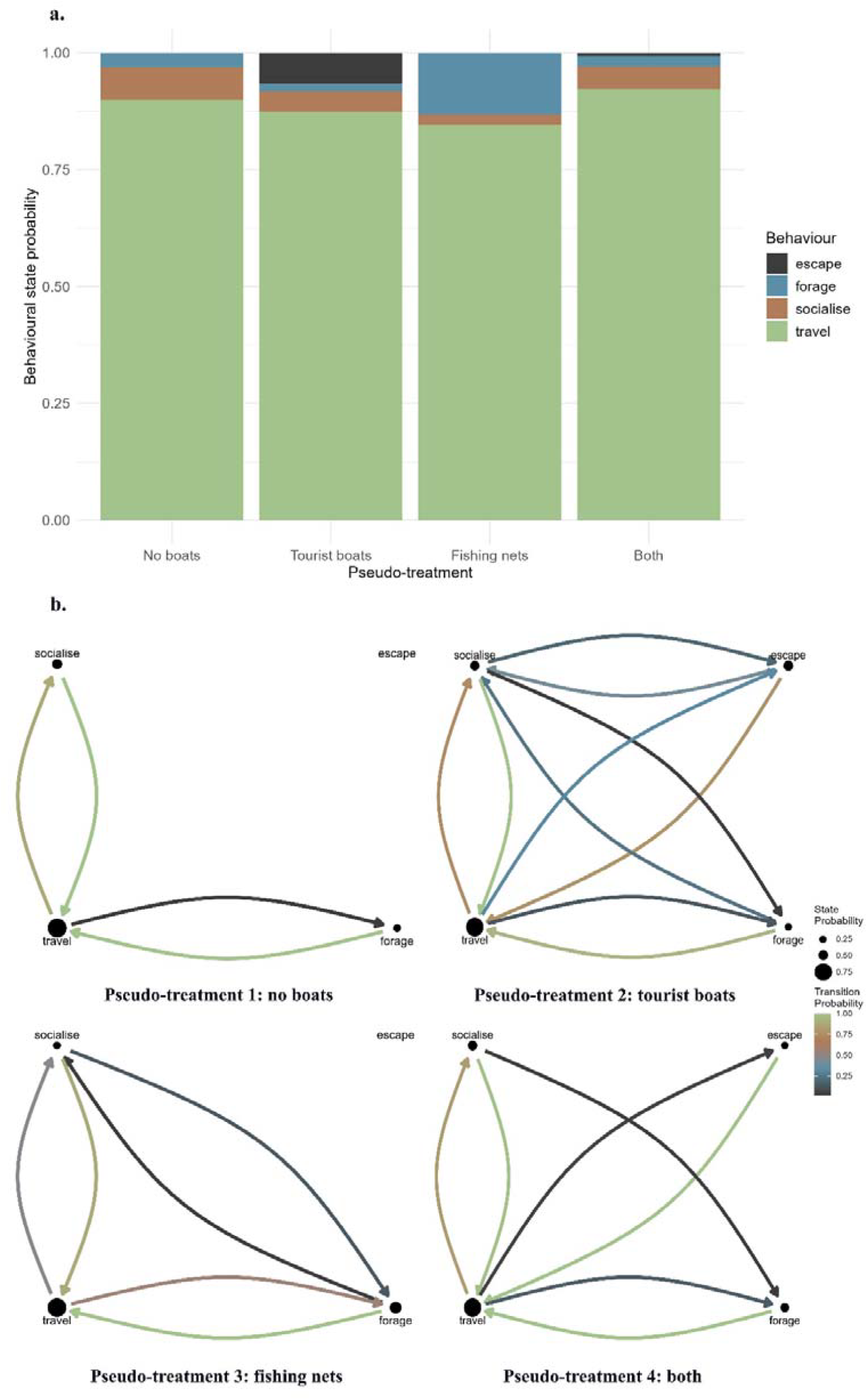
a) Baseline probabilities of different behavioural states in the four pseudo-treatments, b) transition probabilities across different behavioural states in the four pseudo-treatments

The probability of transitioning from one behavioural state to another varied across the four pseudo-treatments and most behavioural transitions occurred from travelling state (Figure 3). In the case of no disturbance (“no boats”), travelling groups mostly switched to socialising (probability = 0.89). When tourist boats were present (“tourist boats”), the switch to socialising decreased by 30% and transitions to escape were observed. Groups near fishing nets (“fishing nets”) were nearly equally likely to switch to foraging (0.55) or socialising (0.45) from travelling. However, even in this case, groups were less likely to switch to foraging (by 75%) when tourist boats arrived (“both”). Escapes only occurred when tourist boats were present (“tourist boats”), but groups were 88% less likely to escape and 55% more likely to forage when fishing nets were also present.

### 3.2. Modulating effects of dolphin behaviour

Behavioural states and transitions were governed by similar factors with some key differences. When only tourist boats were present, dolphin groups were more likely to escape as boats approached closer and as more boats surrounded them. Increasing numbers of boats also decreased the probability of switching from travelling to socialising or foraging. Around fishing nets, groups were also more likely to switch to foraging when they moved closer to nets and to socialising when away. While these transitions decreased in the low-catch season, groups foraged for longer durations after switching. When both boats and nets were present, none of the variables could explain transitions in behavioural states in dolphin groups. However, groups were more likely to escape as tourist boats approached closer or were present in greater numbers (Figure S3, Supplementary Material S1). Similarly, groups continued to forage near fishing nets and escaped when further away in presence of tourist boats.

Group size had significant impact on dolphin behavioural state transitions (Figure 4). Switches to foraging were more common in smaller groups in all cases. Smaller groups were also more likely to escape by diving in the presence of tourist vessels (“tourist boats” and “both”) while larger groups swam away from boats instead. When neighbouring groups were present, focal groups of all sizes were more likely to switch to foraging or socialising.

**Figure 4:.**
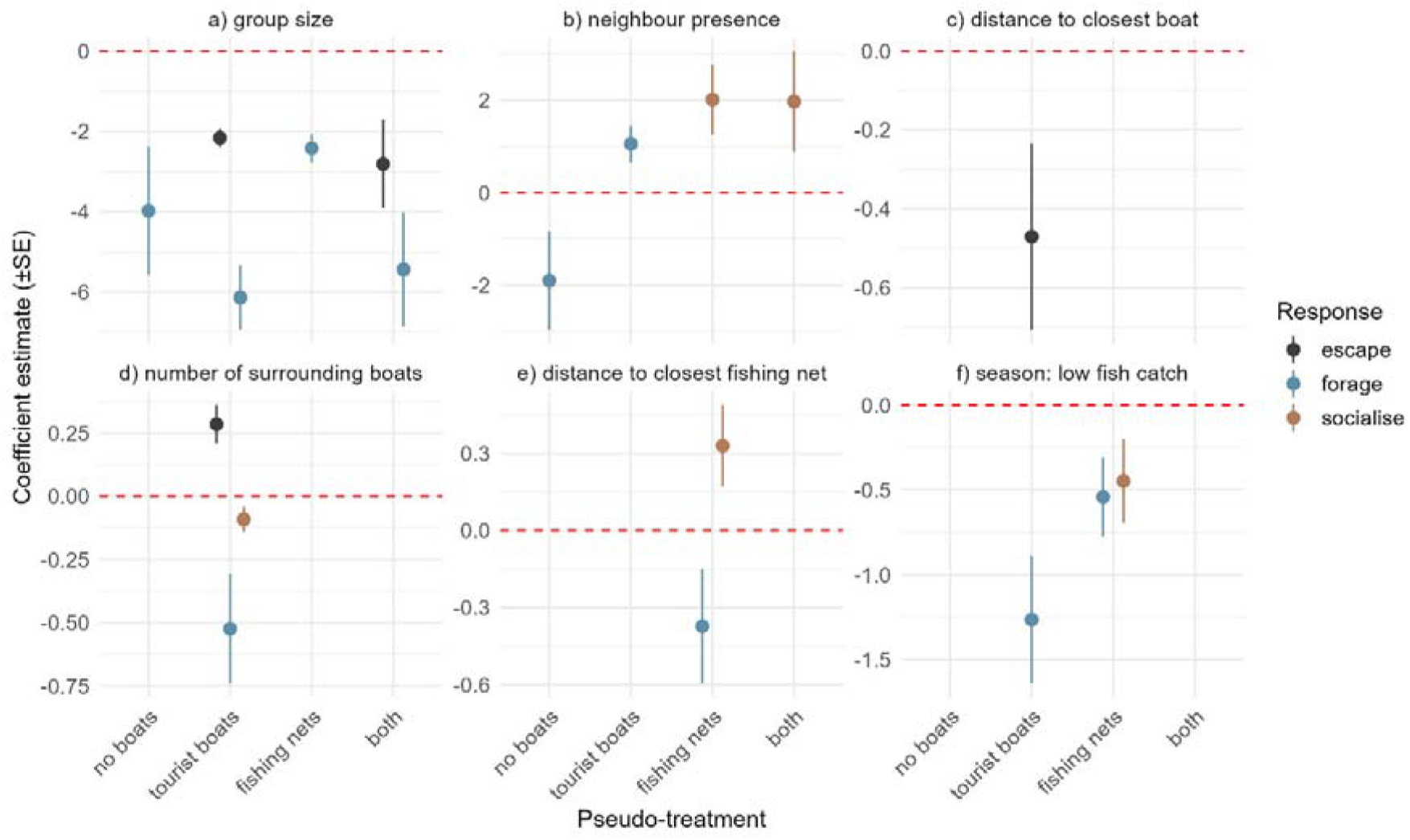
Coefficient estimates for different predictors (a to f) driving the transition from the behavioural state of travel to other states for the four pseudo-treatments

## 4. Discussion

We found that fishery and tourism activities altered behavioural activity budgets and introduced novel behavioural transitions in coastal dolphins. Specifically, Indian Ocean humpback dolphins only expressed escape behaviours when approached by tourist boats and foraged close to (and maybe from) fishing nets, strengthening observations on other dolphin species (Piwetz, Lundquist & Würsig, 2015; Cecchetti et al., 2017). Smaller groups, comprising 1-4 individuals, changed behavioural states more frequently in response to such interactions with human activities, especially when fish-catch availability declined. This study thus provides broad insights for managing coastal human activities and decreasing their threats on threatened delphinids.

Dolphin group responses to tourist boats varied by group size and intensity of pursuit. Groups could be chased by up to 20 tourist vessels at a time and group cohesion increased with approaching boats. As the chase progressed, individuals spent more time on the surface likely due to exhaustion, which in turn increased chances of injury from propeller strikes. Groups were not more likely to split or merge with each other in such conditions; however, in some cases, we recorded small groups diving synchronously and resurfacing as two subgroups at different intervals at different spots. In cases where groups had split, the tourist boats diverted towards one subgroup as the other escaped and splitting perhaps serving as an avoidance strategy. Larger groups may help buffer these effects; while large groups also moved away from tourist vessels, they did not necessarily escape by diving, which could be due to increased vigilance (Filby, Stockin & Scarpaci, 2014) or coordination challenges (Klarevas-Irby, Nyaguthii & Farine 2025). Further research could model the implications of long chases on dolphin energetics and survival, as well as investigating why larger groups appear less sensitive.

Smaller groups were often seen synchronising their dives towards fishing nets and surface away in different directions, suggesting they may be taking fish from or even herding fish into the net. Dolphin groups have often been recorded herding fish into nets that act as physical barriers for the fish (Hersh et al., 2025) and sometimes this may benefit both dolphins and fishers (e.g., Louzada, 2013). In contrast, larger groups also engaged less with fishing nets; such groups may be more efficient at hunting fish collectively (Benoit-Bird & Au, 2009), and their co-occurrence with fishing nets could simply be due to mutual attraction to fish in the area. In general, engagement with fishing nets also increased with declining fish-catch availability; dolphins were more likely to forage near/from fishing nets and for long durations, when fish-catch was low. We could not verify if foraging rates were different across gillnets or purse seines, but increased interactions in April/May with fisheries coincide with increased dolphin mortality in Goa (MMRCNI, 2025).

In cases where fisheries and tourism co-occurred, dolphin responses to both were intermediate but more complex. Dolphins were less likely to forage as tourist vessels disturbed them, but equally less likely to escape, likely because fish caught in (or near) fishing nets kept them engaged. This reflects a trade-off between risk and resources where small dolphin groups may be maximising their foraging gains (Lin et al., 2020) despite risk from a perceived threat such as an approaching tourist boat. Interestingly, dolphins displayed escape responses even to few (1-2) or distant tourist boats (over 10 meters away) when fisheries and tourism co-occurred.

The interactions between dolphins and humans emphasize the importance of shared coastal habitat for both, rendering simplistic solutions like removing human activity to conserve endangered species difficult. Unregulated tourism can severely disrupt dolphin activity (reduced foraging), driving long-term effects on their survival (Sumanapala & Wolf 2024). Simple rules, like limiting the number of tourist vessels and their approach distance to dolphins, especially in case of small groups, can help mitigate these impacts without compromising livelihoods (Zielinski et al., 2022). Similarly, compensations schemes for damaged nets, and for releasing dolphins trapped in nets can help reduce conflict with fisheries if implemented well (La Manna et al., 2024). When dolphins are foraging near nets, even a few tourist vessels can decrease their foraging activity, and so tourist vessels could be instructed to refrain from approaching dolphins at these times. While these management actions are necessary, other factors such as underlying fish availability for both dolphins and fishers can drive negative interactions between the two (Cantor, Farine & Daura-Jorge 2023). Thus, a holistic, ecosystem-based approach is required to improve fish abundance by regulating unsustainable and/or illegal fisheries while also improving habitat quality (Stobutzki et al., 2006).

By combining technology (drones), a novel tracking algorithm and an analytical framework for examining behavioural transitions, we were able to assess when and how dolphins respond to multiple human activities. We not only objectively quantified behaviour by measuring parameters such as speed and direction, but also gathered data on multiple groups simultaneously, comparing both differences in behavioural states across pseudo-treatments as well as how and when behavioural changes occurred. All these pose significant advancements in the field of behavioural ecology (Hughey et al., 2018), especially for studying elusive species like cetaceans.

## Data availability statement

All data and codes relevant to this study will be published as open access in the GitHub repository of IS upon editorial decision.

## Acknowledgements

This paper on the PhD research of IS at CES, IISc which was supported by the Prime Minister’s Research Fellowship. We thank Mrs P. Mitra, the Drishti Foundation, the Sinquerim dolphin watching association and the fisher community of Goa for their support. IS would like to thank Prof P. Berggren for his help when developing this work. Research permits were obtained from the Goa forest department and the Flag Officer Commanding, Indian Navy, Goa. We thank the Rufford Foundation for funding support through their small grants program, and the Dakshin Foundation for funding, logistical support, and providing ethical clearances.

