## Supplementary Material S1 for "Human activities drive novel behaviours and transitions in dolphins"

Extracting data, geo-referencing, and combining video data

To obtain data of dolphin groups, boats, and fishing nets from high-resolution videos, we followed the methodology outlined by Samad et al. (2025). The Convolutional Neural Network (CNN) model trained by the authors was used to identify individual dolphins in each frame of every video. The identities of individual dolphins were connected using a Hungarian algorithm that combines unique identifiers in two consecutive frames based on the overlap between the predicted and the expected position of an animal in the following frame (Hamuda et al., 2018). Such trajectories could only be obtained for individuals on the surface and were lost when dolphins dived. To minimize false positive detections, we only included identities that were tracked for at least 2 seconds. Since our videos were recorded from heights between 60-100 meters above sea level, dolphins were often seen very close to each other. Therefore, in cases when individuals moved close to one another, their unique identifiers were either lost or exchanged, because of which were not able to consistently track all individuals in a group. To overcome this challenge, we assigned group identifiers using a clustering algorithm, and estimated group trajectories instead. Group size was estimated by processing detections across multiple frames (up to 4s) to ensure that individuals that were not identified at any one instance were also included in the estimation. Following this, we manually connected trajectories of diving groups by looking at instances of when they dived and resurfaced. Instances where single individuals or small groups resurfaced for a short time (<2 s) and/or were not identified by our CNN model were manually recorded by clicking on the dolphins upon surfacing. We also manually identified and recorded instances of groups splitting or merging with other groups. We used the same protocol to obtain trajectories of different boats on the water surface by processing our videos using the YOLO v8 model (Varghese & Sambath, 2024).

Finally, to estimate the position and structure of fishing nets deployed by fishing vessels, we manually marked the boundary of the fishing net by clicking on its buoys seen on the water surface. We did this at regular intervals of up to five minutes to ensure that we confidently tracked the shape of the net to estimate variables such as the distance between dolphin groups and the net.

The pixel-coordinates for dolphin and boat trajectories and for fishing nets were geo-referenced and translated into real world space for further analysis. Once complete trajectories were obtained, dolphin group movement speed and direction were estimated by averaging the speed and direction of movement of identified individuals at every second. However, since our drone was not always stationary, our estimates were inaccurate at many instances (e.g., unexpectedly high speed estimated at any one second). To overcome this, we used a moving window of the last five seconds to smoothen estimates for every second. All these analyses were performed using and modifying the scripts provided by Samad et al. (2025).

Once independent datasets for dolphin/boat trajectories and net structures were obtained, they were synchronized using timestamps common across all datasets. For every second, we extracted variables including group size, presence of neighbouring groups, distance to the closest tourist vessel, the number of surrounding tourist vessels, and the distance to closest fishing net.

Table S1: A breakup of sampling effort across the study period

| Month | Sampling effort (days) | Drone footage recorded (hrs) |
| --- | --- | --- |
| October, 2023 | 15 | 12.8 |
| November, 2023 | 24 | 12.4 |
| December, 2023 | 14 | 6.0 |
| January, 2024 | 20 | 5.8 |
| February, 2024 | 19 | 2.5 |
| March, 2024 | 23 | 0.4 |
| April, 2024 | 3 | 0.4 |

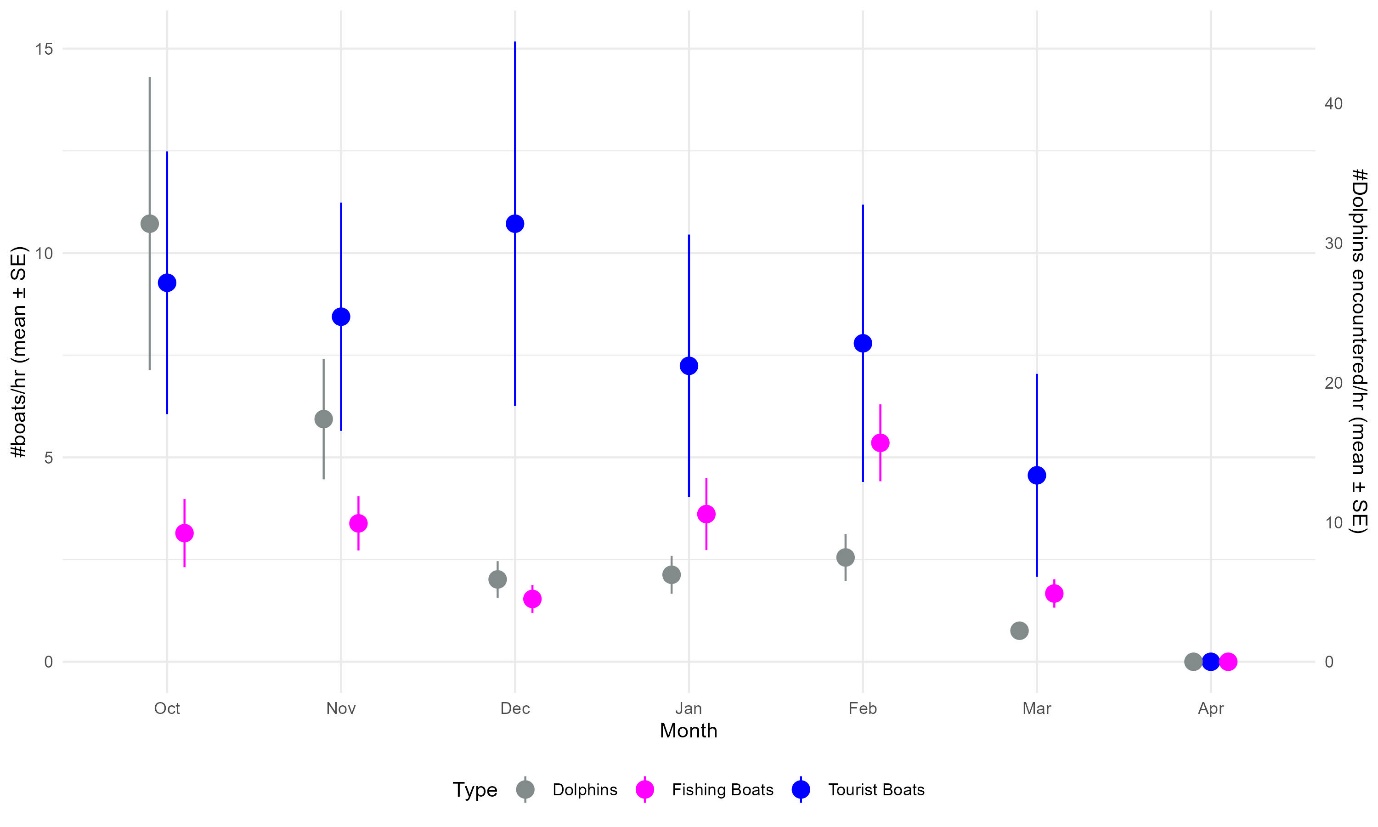

Figure S1: Trends in encounter rates for fishing boats, tourist boats, and dolphins across months

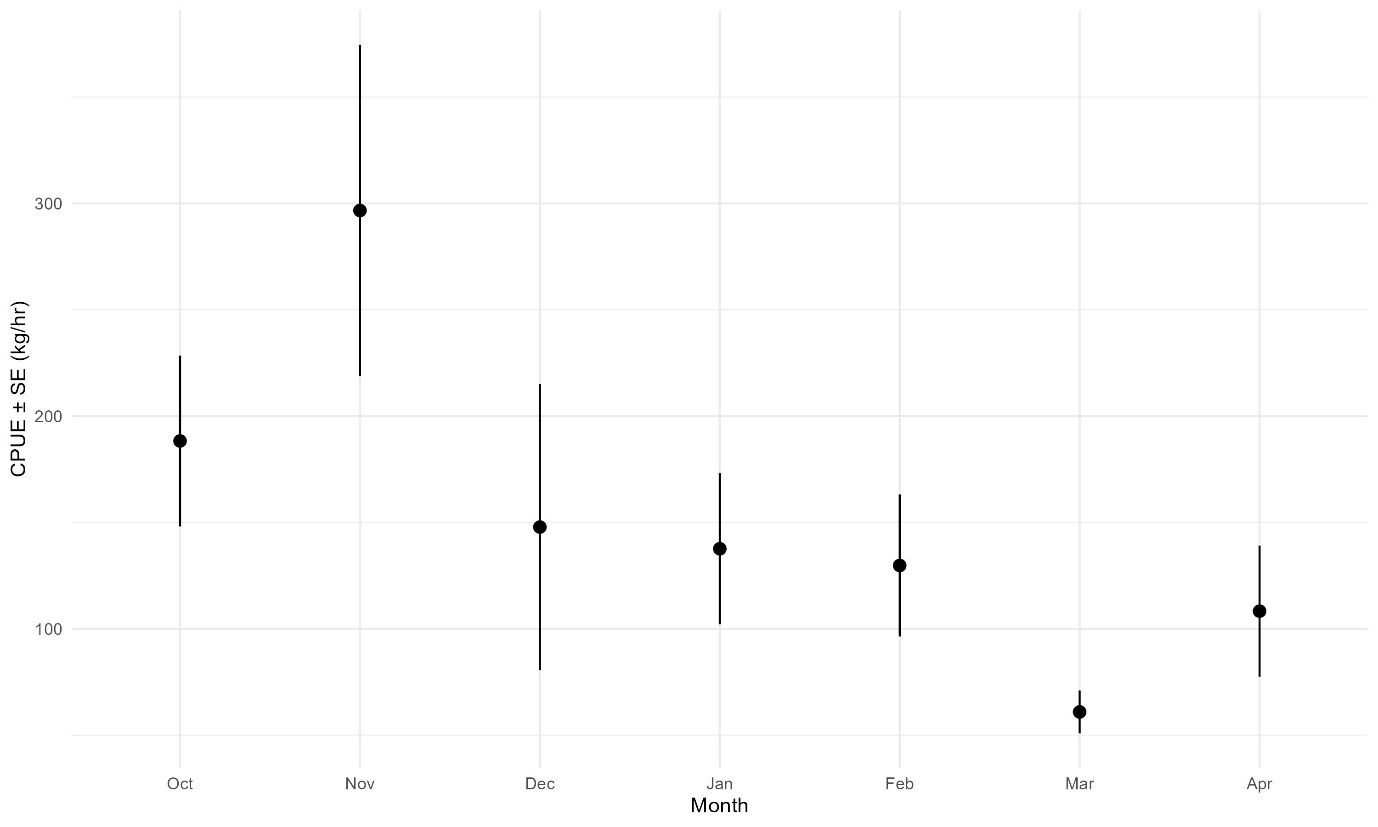

Figure S2: Trends in CPUE across months

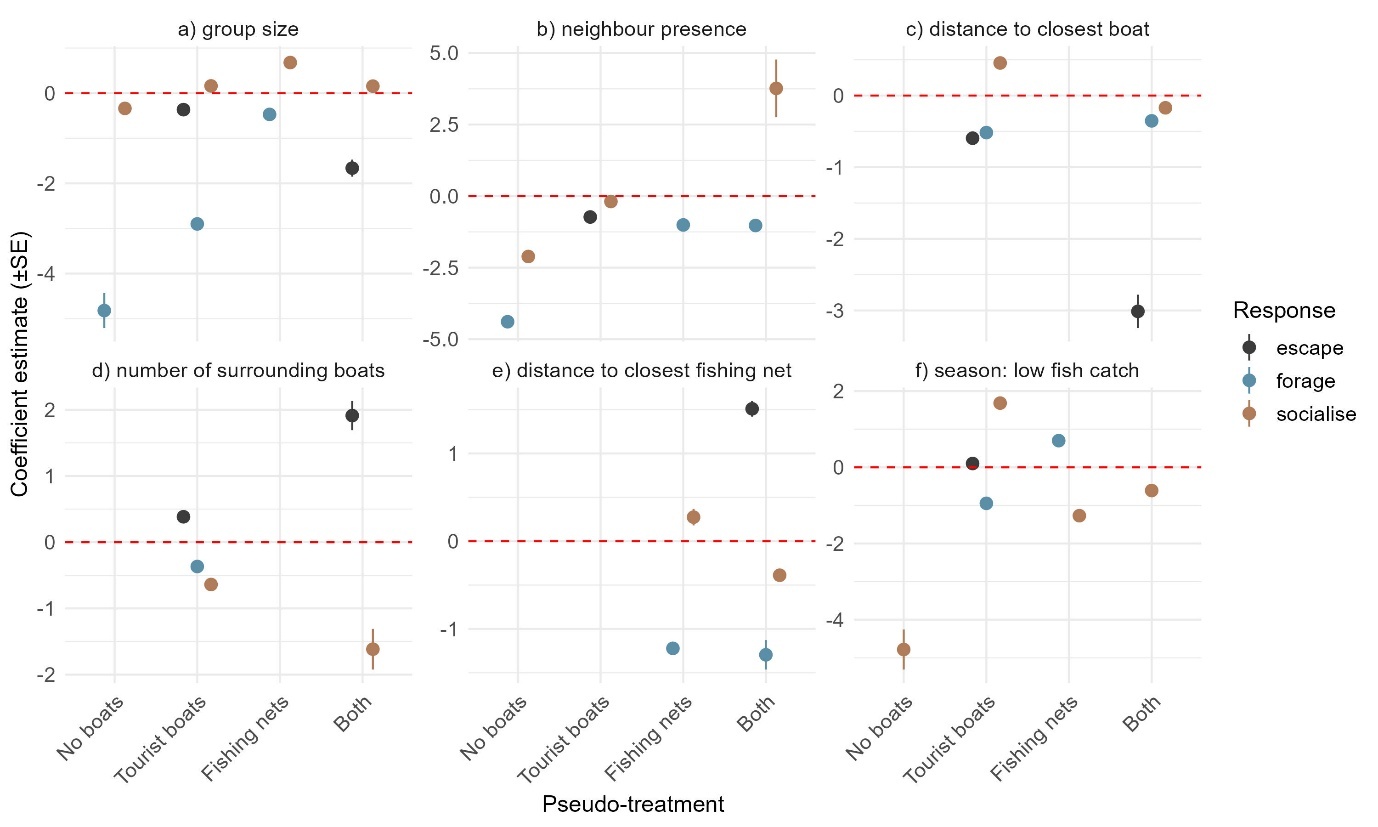

Figure S3: Values of coefficients for different the variables (a to f) allowing the persistence of different behavioural states for the four pseudo-treatments
